# Deployment of tethered gene drive for confined suppression in continuous space requires avoiding drive wave interference

**DOI:** 10.1101/2024.06.24.600398

**Authors:** Ruobing Feng, Jackson Champer

## Abstract

Gene drives have great potential for suppression of pest populations and removal of exotic invasive species. CRISPR homing suppression drive is a powerful but unconfined drive, posing risks of uncontrolled spread. Thus, developing methods for confining a gene drive is of great significance. Tethered drive combines a confined system such as Toxin-Antidote Recessive Embryo (TARE) drive with a strong drive such as a homing suppression drive. It can prevent the homing drive from spreading beyond the confined drive and can be constructed readily, giving it good prospects for future development. However, we have found that care must be taken when deploying tethered drive systems in some scenarios. Simulations of tethered drive in a panmictic population model reveal that successful deployment requires a proper release ratio between the two components, tailored to prevent the suppression drive from eliminating the confined system before it has the chance to spread. Spatial models where the population moves over a one-dimensional landscape display a more serious phenomenon of drive wave interference between the two tethered drive components. If the faster suppression drive wave catches up to the confined drive wave, success is still possible, but it is dependent on drive performance and ecological parameters. Two-dimensional simulations further restrict the parameter range for drive success. Thus, careful consideration must be given to drive performance and ecological conditions, as well as specific release proposals for potential application of tethered drive systems.

## Introduction

As Darwin said, “Natural selection is daily and hourly scrutinizing, rejecting that which is bad, preserving and adding up all that is good.” This principle also applies when humans engineer organisms for specific purposes. Traits introduced through genetic engineering must be capable of persisting under natural selection. Therefore, to ensure the survival and proliferation of these traits, despite potential fitness costs, it is crucial to develop genetic constructs that can increase in frequency within populations (Burt and Trivers, 2006; Gould, 2008).

Gene drives are selfish genetic elements that meet this criterion, able to spread quickly through super-Mendelian inheritance, which equips them with the ability to help control a population after an initial release of transgenic organisms (Esvelt et al., 2014). This efficiency is critical given the escalating threats from pathogens, pests, and invasive species, which now demand innovative solutions (Alphey, 2014; Burt, 2014; Champer et al., 2016; Esvelt et al., 2014). Mosquito-borne pathogens such as malaria, dengue virus, Zika virus, chikungunya virus, yellow fever virus, and West Nile virus pose serious threats to human health (Gubler, 2011; Tahir et al., 2019; Weaver and Reisen, 2010). Additionally, agricultural pests compromise productivity and food safety, resulting in costs of approximately $540 billion annually if left unchecked (Willis, 2017). Moreover, invasive species are increasingly damaging ecosystems, affecting fields, fisheries, and forests worldwide (Teem et al., 2020). Given these challenges, gene drives offer a promising solution. These techniques provide a more effective, controllable, economically friendly, and environmentally sustainable approach to managing populations of harmful pathogens, pests, and invasive species (Esvelt and Gemmell, 2017).

The capability of gene drives to spread rapidly stems from their ability to bias their own inheritance, either by duplicating themselves or by eliminating a competing allele (Basgall et al., 2018; Burt and Crisanti, 2018; Roggenkamp et al., 2018). Different gene drive designs have different propagating capabilities and can be divided into three categories: “invasive”, “confined” and “self-limiting” (Wang et al., 2022b). Invasive drives can spread rapidly but uncontrollably with low initial release requirements. These include homing drives, which encode an endonuclease such as Cas9 that cleaves a target site in the homologous chromosome and then converts the wild-type allele into a drive allele during homology-directed repair (Deredec et al., 2008). Confined drives spread more slowly and tend to remain in a local population due to non-zero introduction thresholds. They include underdominance drives in which drive/wild-type heterozygotes are less fit than either drive or wild-type homozygotes. The introduction threshold forms because both drive and wild-type alleles can be removed from the population (Wang et al., 2022b). In confined drives, only when the release frequency exceeds the introduction threshold can the drive elevate its own frequency without being eliminated by wild-type. Self-limiting drives, like split drive (Champer et al., 2019), are unable to retain high frequency in the long-term in the presence of wild-type alleles. These systems can still bias inheritance to slow their decay or even temporarily increase, but they will eventually decline if they fail to reach fixation (Wang et al., 2022a). All these drive designs with different propagating capabilities give multiple options to achieve levels of confinement appropriate to address different ecological problems.

Gene drives can affect a population in two ways, by propagating desired traits to the target population as a “modification drive” or by reducing the target population, which is called “suppression drive”. Modification drives with different propagating capabilities can achieve different levels of confinement. Homing drive is a powerful but invasive drive(Adolfi et al., 2020; Gantz et al., 2015), which can be of concern from both a scientific and public relations standpoint (Rode et al., 2020). Confined gene drives can only spread when the release frequency is above the introduction threshold and can only expand between separated populations when the migration level exceeds a separate threshold (Champer et al., 2021a). CRISPR toxin-antidote gene drives usually fall into this category and are relatively straightforward to construct because they can utilize end-joining repair after DNA cleavage rather than just homology-directed repair, which may have a shorter window of opportunity (Heyer et al., 2010) and relatively stricter requirements (Lin et al., 2014). One promising design is Toxin-Antidote Recessive Embryo (TARE) drive, which targets an essential and haplosufficient gene with the drive providing the recorded copy of the target gene. This means that individuals with two nonfunctional resistance alleles cannot survive and will be removed from the population, but any individual with a drive or wild-type allele will be viable (Champer et al., 2020b; Oberhofer et al., 2019). TARE drive is fairly powerful but still confined with a low introduction threshold (Champer et al., 2020a). Derived from TARE drive, 2-locus TARE drive is more confined and has a non-zero introduction threshold even without fitness costs. It involves two drive alleles at different genomic loci, with each drive allele targeting the gene for which the other provides rescue. These CRISPR toxin-antidote gene drive variants allow for a variety of thresholds depending on the arrangement of drive components (Champer et al., 2021a).

Unlike modification drive, the realization of successful suppression drive requires not only basic propagating ability, but also strong enough genetic load to suppress the population. With good performance parameters, a homing suppression drive can meet this criterion (Kyrou et al., 2018), but this type of drive could spread to non-target populations. When we aim to maintain control by using a drive type with weaker propagation capabilities, genetic load (referring to the power of the drive to suppress the target population) will often be insufficient. Thus, many confined modification drives cannot be transformed into effective suppression drives, such as TARE (Champer et al., 2020a). Yet, this type of drive design has broad development prospects because its confinement is conducive to its application in the field of sustainably controlling harmful pests and removing invasives in local areas. One possible strategy to achieve confined suppression is using Toxin-Antidote Dominant Embryo (TADE) suppression drive. This system disrupts essential fertility genes and also targets a haplolethal gene using Cas9, while the drive allele provides a rescue copy of the haplolethal gene (Champer et al., 2021a; Champer et al., 2020a; Zhang and Champer, 2024; Zhu and Champer, 2023). However, the haplolethal target gene makes these drives challenging to engineer.

One possible way to achieve confined suppression is to split the confinement and the suppression aspects into two separate drives and then “tether” them together (Dhole et al., 2019). Tethered gene drives combine a complete confined modification drive and an incomplete homing suppression drive. The latter lacks a component needed for drive, but this is provided by the confined drive. Thus, the homing suppression drive can only spread in the region occupied by the confined drive. A tethered homing drive with a TARE drive that provided Cas9 was recently confirmed to be experimentally feasible (Metzloff et al., 2022).

While promising, modeling for tethered drives has been limited to a few representative examples in panmictic populations (Dhole et al., 2019). Although panmictic models can help identify key parameters that facilitate the spread of gene drives throughout populations and show dynamics for small populations or a widespread release (Deredec et al., 2008), it has become evident that a comprehensive understanding of gene drive outcomes in many scenarios requires consideration of spatial dynamics. Spatial structure may lead to an unstable coexistence between drive and wild-type alleles, in contrast to panmictic models which often predict dominance of one over the other or a stable equilibrium between them. Previous research on suppression drives revealed outcomes where regions were either empty, occupied solely by wild-type individuals, or containing a mix of wild-type and drive alleles (Bull et al., 2019; North et al., 2020). Other research highlighted that this can trigger “chasing” dynamics where wild-type individuals repopulate areas that the drive had previously been cleared of individuals by the drive, which may contribute to more dynamic outcomes and unstable persistence (Champer et al., 2021b). With tethered drives and their greater number of genotypes (due to the presence of two drives), new types of behavior may be found that are not seen in idealized panmictic models, potentially drastically changing the final outcome of a gene drive release.

In this study, we investigated the outcomes of tethered gene drives using a population model that incorporates spatial continuity. Based on our simulations, continuous space significantly influences outcomes. We found variable wave interference among the three group of wild-type individuals, TARE drive individuals, and homing suppression drive individuals. As the TARE drive wave continues to advance and the homing drive wave catches up, ideally, the two drive waves would continue to advance and clear out an empty area behind them, achieving suppression. However, the homing drive may instead weaken the TARE drive wave, leading to TARE drive loss or providing an opportunity for non-drive individuals at the wave front to escape. When these non-drive individuals penetrate through the drive waves, they become founders in empty space, leading to chasing dynamics and complex outcomes. Our simulations illustrate all these results and demonstrate the outcomes of tethered drives under various models and parameters, showing which scenarios should be avoided and thus laying the groundwork for development of release plans for future applications.

## Methods

### Tethered homing drive design

Tethered drives are composed of two genetic elements, confined modification and suppression, which are placed in different loci. Two tethered homing drive designs based on TARE or 2-locus TARE are tested. Both have an incomplete homing suppression drive, which is placed inside an essential but haplosufficient female fertility gene and lacks the Cas9 element. Only when an individual has a TARE drive allele (with Cas9) can the homing suppression allele conduct homology-directed repair and germline drive conversion. Drive homozygous females (or females with any combination of drive alleles and nonfunctional resistance alleles) are sterile. The confined modification drive has two different variants. The first is a TARE drive placed inside in a haplosufficient gene, for which it provides rescue. The drive contains Cas9. If an organism has two nonfunctional alleles of the haplosufficient gene, it will be nonviable. The second drive type is 2-locus TARE drive, where two genetic constructs are each placed inside different haplosufficient genes (for which they provide rescue) with a reciprocal targeting. Each of the loci also contains Cas9. Due to the removal of drive alleles by mutual disruption, it has a modest introduction threshold of 18% in the absence of fitness costs.

In our simulations, offspring inherit alleles from their parents at random, which can be affected by drive activity. For the TARE drive, if an offspring inherits a wild-type allele from a parent who also carries a drive allele, this allele has a chance of being converted into a nonfunctional resistance allele, determined by the germline cut rate. Moreover, if the mother carries any drive alleles, any remaining wild-type alleles in her offspring may also be converted to nonfunctional alleles at a probability equal to the embryo cut rate. For the homing drive, only when an offspring has Cas9 from a TARE drive allele can any drive activity take place. If an individual would normally receive a wild-type allele (at the homing site) from a parent who possesses a homing drive allele and a TARE allele, then this wild-type allele may transform into a drive allele, determined by drive conversion rate. It may also have a chance to become a nonfunctional resistance allele, determined by the resistance allele formation rate. If the mother carries at least one homing drive allele (and a TARE allele), then the embryo cut rate is the fraction of wild-type alleles remaining in the offspring that will be converted to nonfunctional resistance alleles.

By default, TARE drive was set with a germline cut rate of 1.00 and an embryo cut rate of 1.00, and homing suppression drive was set with a drive conversion rate of 0.95 (remaining alleles in the germline are converted to resistance alleles) and an embryo cut rate of 0.15. To get the genotype-based fitness, we use multiplicative drive allele fitness for TARE, assuming that each allele has an independent fitness cost. For homing suppression drive, we assume that fitness costs only apply to female drive/wild-type heterozygotes. By default, there are no fitness costs (fitness is set to 1.0).

### Simulations

All simulations were implemented in the forward-in-time genetic simulation framework SLiM, version 4.0 (Haller and Messer, 2023). The model was based on previous studies of TARE drive (Champer et al., 2020a) and homing suppression drive (Champer et al., 2021b). All simulated data, SLiM simulation models, and data-processing scripts are available on GitHub (https://github.com/jchamper/ChamperLab/tree/main/Tethered-drive).

### Panmictic population model

Simulations were first implemented in a panmictic population with the following reproduction and survival rules: (1) Generations began with mate choice. Each fertile female selects a random potential male, which is then confirmed with a probability equal to its fitness. If not confirmed, the female will select another potential mate and can make up to 20 attempts. If no male is selected or none is available, the female does not reproduce. (2) Female fecundity is determined based on a Beverton–Holt curve as 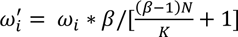, where 𝜔*_i_* is the baseline fecundity due to genotype, *N* is the total population size, *K* is the population carrying capacity, and 𝛽 is the low-density growth rate, with a default value of 10. (3) The actual number of offspring is generated based on a binomial distribution with 𝑛 = 50 and 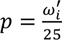, yielding a maximum of 50 offspring and an average of two offspring for a female with fitness 𝜔_i_ = 1 at normal population density. (4) Offspring genotypes are obtained according to the tethered drive mechanisms.

Simulations were initialized with a total population of 𝐾 = 50,000 with different drive release methods. The first release mode was separate 1-genotype release, referring to releasing one group of TARE drive homozygotes (**D_T_D_T_+_H_+_H_**) (Table 1) and another group of homing suppression drive heterozygotes without the TARE drive (**+_T_+_T_ D_H_+_H_**). The second release mode involved releasing one group of TARE drive homozygotes (**D_T_D_T_+_H_+_H_**) and another group of individuals that were TARE drive homozygotes and homing suppression drive heterozygotes (**D_T_D_T_D_H_+_H_**). The release ratio was defined as relative to the wild-type population size.

**Table 1.** Genotype abbreviations.

| Abbreviation | Meaning |
| --- | --- |
| **D<sub>T</sub>** | TARE drive allele |
| **+T** | TARE wild-type allele |
| **D<sub>H</sub>** | Homing suppression drive allele |
| **+H** | Homing suppression wild-type allele |

### One-dimensional spatial model

The panmictic model was extended into a one-dimensional continuous space model with a length of 1. The one-dimensional arena was set to have reprising boundaries, which means that when an individual’s position would fall outside the arena, a new position was drawn instead. In the 1D spatial model, reproduction and interaction rules were changed. We set 𝑑𝑖𝑠𝑝𝑒𝑟𝑠𝑎𝑙 𝑑𝑖𝑠𝑡𝑎𝑛𝑐𝑒 to represent individual’s mobility and interactions. (1) When choosing a mate, females could only choose mates within a distance equal to the 𝑑𝑖𝑠𝑝𝑒𝑟𝑠𝑎𝑙 𝑑𝑖𝑠𝑡𝑎𝑛𝑐𝑒. (2) Local competition was considered, so the fecundity of female was 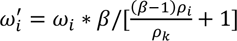, We define the expected carrying density interaction strength as 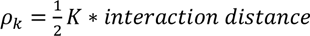, where *K* is the population carrying density (and total capacity in the arena of length 1). Next, we define a local density, 𝜌_i_, which represents the local interaction strength for each female within the specified interaction distance, set at 0.01. Each individual contributes a density strength of “1” if they occupy the same position as the focal female. This strength linearly declines to zero as the distance approaches 0.01. (3) Offspring were distributed around the mother. The distance between offspring and their mother was drawn from a normal distribution with zero mean and standard deviation equal to 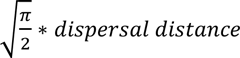, producing an average displacement equal to 𝑑𝑖𝑠𝑝𝑒𝑟𝑠𝑎𝑙 𝑑𝑖𝑠𝑡𝑎𝑛𝑐𝑒.

Simulations were initialized with a total population of 𝐾 = 10,000 and a fixed release ratio. 90% of individuals in the leftmost 5% of the arena were transformed into TARE drive homozygotes (double homozygotes for the 2-locus TARE), and several generations later, 50% of individuals in the leftmost 5% region were transformed into individuals that were TARE drive homozygotes and homing suppression drive heterozygotes. This time was determined based on the drive wave speeds, such that the homing drive wave of advance would meet the TARE drive wave in the middle of the arena. Specifically, timing was adjusted so that in the middle 10% of the area, both drives, would reach 50% allele frequency at the same time, on average.

### Two-dimensional spatial model

The one-dimensional model was extended into two-dimensional space (with a 1 x 1 arena) to conduct simulations more representative of a natural landscape. We set 𝑑𝑖𝑠𝑝𝑒𝑟𝑠𝑎𝑙 𝑑𝑖𝑠𝑡𝑎𝑛𝑐𝑒 to represent an individual’s mobility. In the 2D spatial model, both the horizontal and vertical distance between offspring and their mother were separately drawn from a normal distribution with zero mean and standard deviation equal to 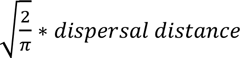, yielding an integrated average displacement equal to 𝑑𝑖𝑠𝑝𝑒𝑟𝑠𝑎𝑙 𝑑𝑖𝑠𝑡𝑎𝑛𝑐𝑒.

Simulations were initialized with a total population of 𝐾 = 50,000 with single 2-genotype release mode and a fixed release ratio. 90% of individuals within a circle of radius 0.1 at the center were transformed into TARE drive homozygotes (double homozygotes for the 2-locus TARE), and several generations later, 50% of individuals in a circle of radius 0.1 were transformed into individuals that were homozygous for TARE and heterozygous for the homing suppression drive. The second release time was calculated depending on the drive wave speed to ensure that the drive waves met (reached 50% allele frequency at the same time) at a radius between 0.225 and 0.275.

### Penetration detection

A penetration event is considered indicative of a potential scenario where wild-type alleles pass through the drive wave, representing an instance where wild-type has escaped to potentially form a new wave moving away from drive individuals. This potentially triggers a chasing event. Our one-dimensional arena was split into 100 segments, allowing for precise tracking of allele frequencies and population sizes at each segment. The allele frequencies across segments were subjected to a smoothing process to reduce random noise and facilitate analysis of spatial patterns and trends. Smoothing was performed by recalculating the value of each segment as the weighted average of itself and the three closest segments on each side, with weights assigned based on distance from the central segment: 100% for the segment itself, 75% for the first adjacent segments, 37.5% for the second, and 18.75% for the third. By analyzing the smoothed allele frequency distribution curves for each generation, we identified the positions of local peaks, which we define as higher than the five values preceding and following them, and troughs, which are lower than the five values preceding and following them. These are classified into three categories: peaks of the drive allele, peaks of the wild-type allele, and troughs representing empty spaces. We considered a potential wave penetration event to have occurred if a pattern where a wild-type peak is flanked by a drive peak on one side and an empty trough on the other is observed for more than three consecutive generations.

## Results

### Characteristics of Tethered homing drives

Our tethered drive system involved a complete TARE drive for confinement and an incomplete homing drive for suppression. The TARE drive sits in an essential and haplosufficient gene and consisted of a toxin (Cas9 and gRNAs targeting this gene) and an antidote (a recorded functional version of the target gene). Drive/wild-type heterozygotes will disrupt the wild-type allele in the germline. Wild-type alleles in the progeny of females will also be converted to nonfunctional resistance alleles though maternal deposition of Cas9 and gRNA at the embryo stage. Because it is an essential and haplosufficient gene, individuals with two copies of nonfunctional resistance alleles will die (Figure 1A), increasing the relative frequency of the drive.

**Figure 1.**
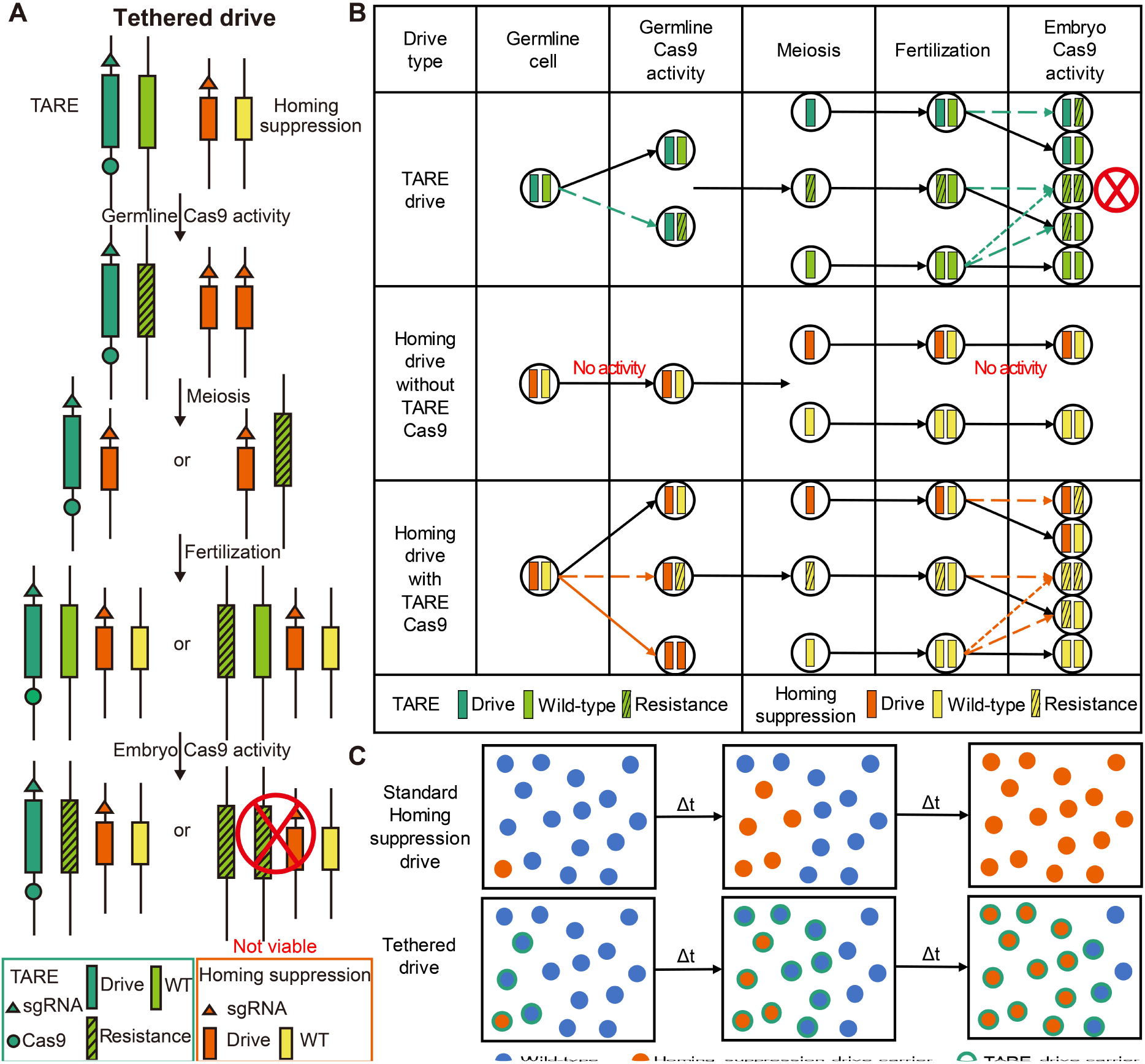
Tethered drive mechanism. (**A**) The tethered homing drive is demonstrated in a female that is heterozygous for a TARE drive allele and a homing suppression drive allele. Germline activity takes place, followed by mating with a wild-type individual and cleavage in the early embryo from maternally deposited Cas9 and gRNA. (**B**) The mechanisms of TARE drive, homing drive without TARE, and homing drive with TARE, from the germline to early embryo. First, in TARE drive, Cas9 activity occurs in the germline of drive-carrying individuals, disrupting wild-type alleles. Then, embryo Cas9 activity occurs in the progeny of females, which can disrupt any surviving wild-type alleles from the mother or new wild-type alleles from the father. Individuals with two nonfunctional resistance alleles are not viable. Second, when homing drive individuals do not have a source of Cas9, homing drive activity cannot occur. Third, in homing drive heterozygotes that also carry the TARE drive with Cas9, Cas9 activity in the germline causes drive conversion by homology-directed repair. Resistance alleles can also be formed in the germline or in the embryo stage from maternal deposition. The colored solid arrows represent drive conversion, the colored dashed arrows show resistance allele formation, and the black solid arrows show normal transitions without Cas9 cleavage. (**C**) The population dynamics of standard homing drive and tethered drive are illustrated by showing three different times after drive release. A standard homing drive, which has no threshold frequency, can spread through a wild-type population after a small release, whereas a tethered homing drive can only spread in TARE-modified population. The TARE drive also has a larger required initial release size.

The incomplete homing suppression drive lacking Cas9 element is located in an essential but haplosufficient female fertility gene, with gRNAs targeting the gene at the drive site. Drive activity can only happen in TARE drive carriers (Figure 1B). Therefore, the homing suppression drive allele is capable of replication solely within the boundaries established by the TARE drive mechanism (Figure 1C), endowing the originally “invasive” homing suppression drive with “confined” characteristics.

Because tethered homing drives represent a relatively flexible system, the nature of their confinement is determined by the modification drive, allowing for the application of various mechanisms to achieve different degrees of confinement. Consequently, a variation of the TARE drive, the 2-locus TARE drive, is also analyzed as part of a tethered drive system. 2-locus TARE drive is composed of two loci, each of which has a TARE drive allele specifically targeting the gene that the other one rescuing (Figure S1), instead of the gene at the same site as the drive. Because the toxin and rescue elements are segregated from each other, there are fewer viable genotypes, causing 2-locus TARE drive to have a significantly higher introduction threshold than the standard TARE drive. A tethered system based on it will therefore have increased confinement (Figure S1).

### Confinement of Tethered homing drives

To evaluate the confinement characteristics of tethered drives, we varied the initial release proportions and observed the outcomes. Because the TARE drive is linked to the homing suppression drive, which can remove TARE drive alleles, the introduction threshold of the confined TARE drive should be higher than when the TARE drive is released alone. Confinement also makes the system distinct from traditional homing suppression drive, which is a zero-threshold system and can quickly spread from even small release sizes. Different variants of TARE drive have different introduction thresholds, which can provide different levels of confinement as needed. Thus, we modeled both TARE-based drive and 2-locus TARE drive.

Two release modes were adopted. The first mode simultaneously released TARE drive homozygotes (**D_T_D_T_+_H_+_H_**) and separately, individuals with only with one copy of homing drive (**+_T_+_T_ D_H_+_H_**). The second release mode had TARE drive homozygotes (**D_T_D_T_+_H_+_H_**) and separately, individuals that were both homozygous for the TARE drive and heterozygous for the homing drive (**D_T_D_T_D_H_+_H_**). 2-locus TARE individuals were homozygous for both TARE alleles. We varied the release frequency of each of the two genotypes independently, relative to the initial population. TARE and 2-locus TARE drives had no fitness costs and 100% germline and embryo cut rates. This should produce a zero introduction threshold for TARE (Champer et al., 2020a) and an 18% threshold for 2-locus TARE (Champer et al., 2021a).

In panmictic models, there were only two major outcomes, either successful population elimination or confined drive loss (Figure S2). When using the separate release method, both confined drives had a higher introduction threshold that confined-only releases, which increased with higher suppression drive release level. This was because the homing drive could eliminate a few confined drive alleles before it could become solidly established, or in some cases, catch up and eliminate the confined drive before it reached the whole population. When releasing individuals with the confined and suppression drive together (in addition to individuals with just the confined drive), because homing carriers themselves also had TARE drive alleles, they could spread more quickly at first, inhibiting the TARE drives more easily. This was counteracted by these individuals possessing the TARE drives in the first place, essentially raising the release amount of this confined drive. The introduction pattern was particularly distinct for the 2-locus TARE drive system, where fewer TARE-only individuals were required for success (Figure 2C-D). However, the total number of released individuals required for success was still increased as the number of 2-locus TARE/homing suppression drive released individuals increased. Even above its introduction threshold, TARE-based tethered drive can have stochastic failure (albeit at a low rate), which was far less common for the 2-locus TARE-based tethered drive, potentially making the latter drive more reliable under certain release scenarios despite its increased introduction threshold. This is likely caused by differences between the endgames of these two drives, leading to a third possible outcome. At high drive frequency, the 2-locus TARE drive can create more nonviable genotypes, making it easier for the drive frequency to increase further. At the same time, with a relatively high initial release the homing drive carrier frequency may catch up to and surpass the TARE drive (Figure S3A). It usually will not be able to pass the frequency of a 2-locus TARE drive. This may lead to a separation between the TARE drive and the homing drive (Figure S3B-C), reducing the suppressive power and resulting in a high equilibrium frequency rather than successful population suppression.

**Figure 2.**
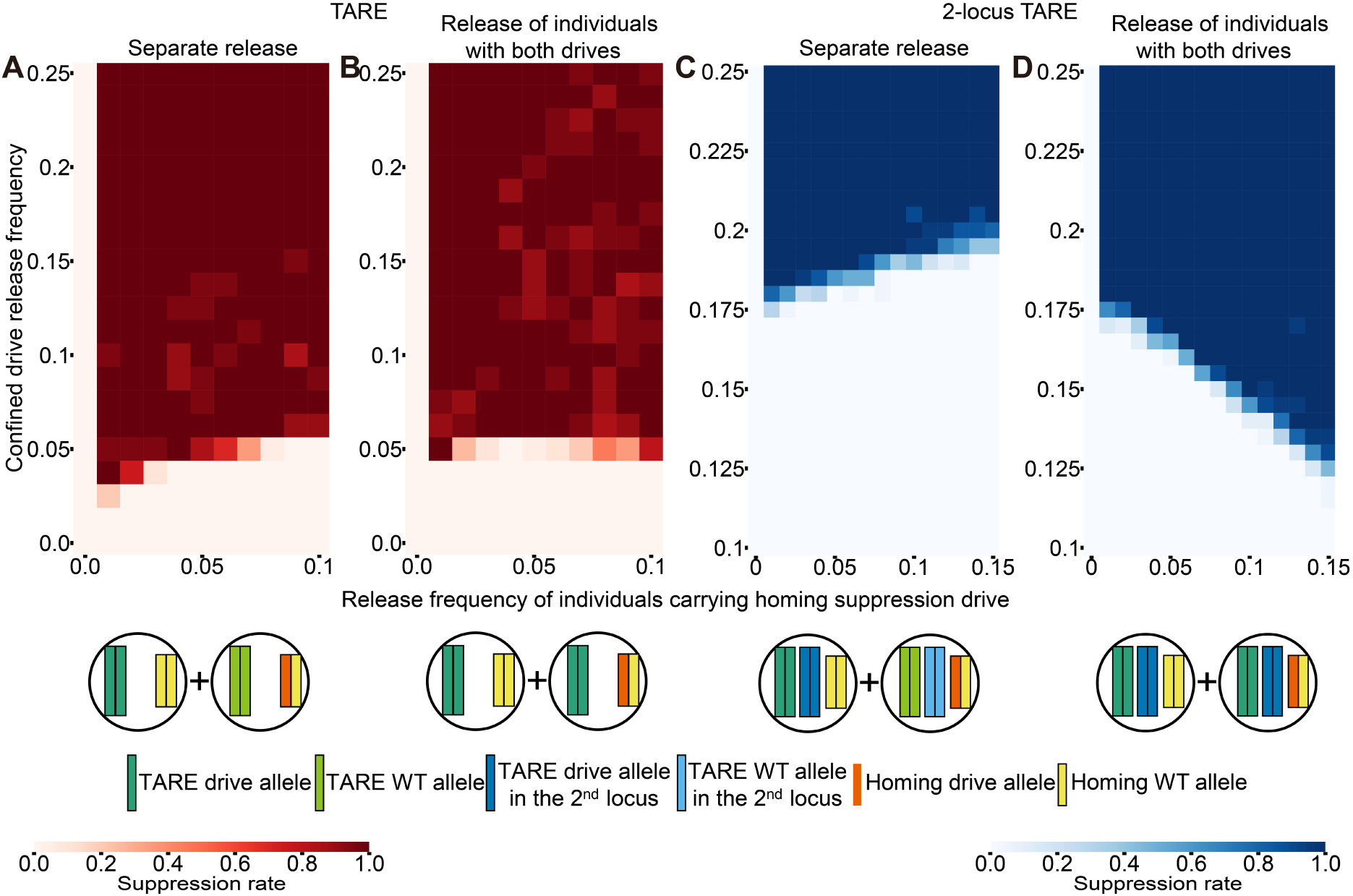
Confinement of tethered drives in panmictic populations. Confined TARE drives and homing suppression drive, both with default parameters, were released at the same time with two different release methods and varying release ratios. The genotypes of the released individuals are shown below the heatmap. (**A-B**) TARE-based tethered homing drives using (**A**) separate releases and (**B**) 2-drive releases. (**C-D**) 2-locus TARE-based tethered homing drives using (**C**) separate releases and (**D**) 2-drive releases.

### Outcomes in the one-dimensional spatial model

While panmictic populations are representative of some real-world situations, individuals in a population will often be spread over a continuous landscape and capable of only moving through a small fraction of that landscape in their lifetimes. To investigate the performance of the tethered drive in such a system, we used a one-dimensional spatial model to allow for simplified study of drive waves. The rationale behind establishing a one-dimensional spatial model lies in the tethered drive’s unique characteristic of exhibiting a “tethered” attribute, where the Cas9-free homing suppression drive relies on the Cas9-based TARE drive. In this system, the homing suppression drive could catch up and undermine the slower TARE drive, removing TARE drive alleles and preventing it from continuing to advance into the wild-type population. To initialize the scenario, the TARE drive was released at one end of the arena, and the homing suppression drive was released later, so that it would catch up to the TARE drive in roughly the middle of the area (Figure 3A).

**Figure 3.**
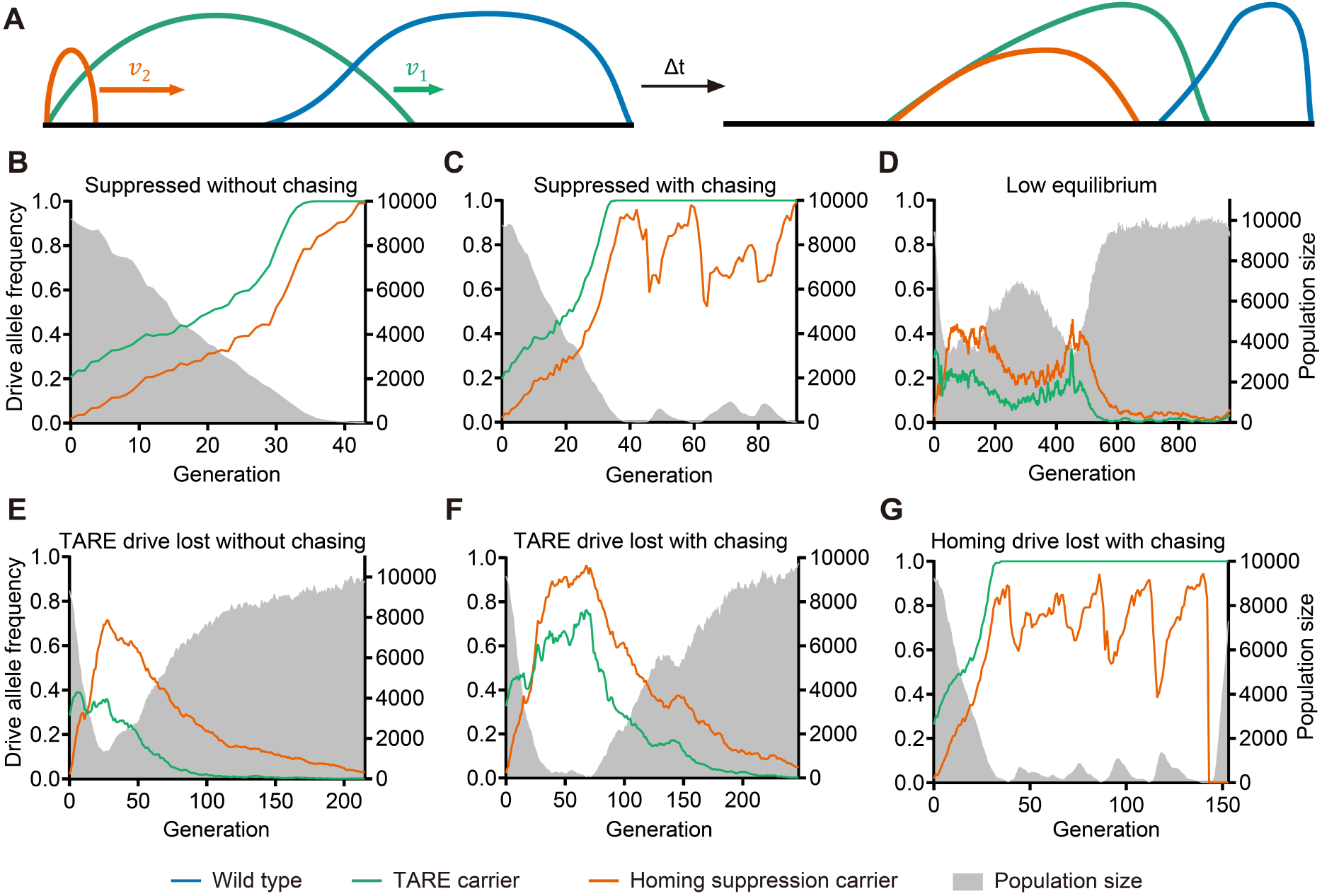
Possible outcomes in tethered homing drives in one-dimensional model. (**A**) The one-dimensional model is shown. TARE drive and homing suppression drive are released sequentially. In general, TARE drive is slower than homing suppression drive, and is undermined by homing suppression drive after it catches up to TARE, contributing to a complex wave interference. Blue, green and orange solid line represents the density of wild-type individuals, TARE drive carriers and homing suppression drive carriers, respectively. (**B-G**) Time-series of different possible outcomes is shown. Green and orange solid line represents the drive allele frequency of TARE drive and homing suppression drive, respectively. The grey shadow represents the population size.

The final outcomes were categorized into successful population elimination, TARE drive lost, homing suppression drive lost, and a situation where both drives remain without eliminating the population. Further, each of these outcomes could occur with and without chasing, in which drive and wild-type alleles persist together with regions of empty space (Champer et al., 2021b). Of these eight outcomes, only six had a moderate to high frequency of occurrence (Figure 3B-G). This includes a “low equilibrium” outcome, which means both drives remained at relatively low frequency for a long time without chasing (this is in contrast to the high equilibrium outcome in panmictic populations, which did not occur in out spatial models). We detected chasing by checking when the wave formed by wild-type individuals penetrated drive-carrying individuals and recolonized an empty area. Usually, unless the TARE drive and homing suppression drive kept balance and continued to advance, chasing almost always occurred, leading to fluctuations in drive frequency (Figure 3D-G).

We also examined the performance of 2-locus TARE to support tethered drive systems in continuous space. However, this version was never successful. The 2-locus TARE wave always collapsed when the homing drive caught up over the full range of parameters.

### Parameters affecting drive in the one-dimensional spatial model

To further clarify the performance of tethered drive in one-dimensional space, homing drive performance parameters were varied, while TARE drive was still assumed to be optimal, due to the lower practical requirements for high efficiency TARE drive (Champer et al., 2020b; Metzloff et al., 2022). In general, successful population elimination was possible over much of the parameter range for varying drive conversion efficiency and embryo cut rate, though usually after a moderate period of chasing of varying length (Figure 4A-C). Only when the homing suppression drive efficiency was over 95% with a relatively low embryo cut rate would the TARE drive be consistently eliminated (Figure 4D-E), followed by the failure of the homing suppression drive due to lack of Cas9. When TARE drive loss was not possible, there was a possibility of the homing drive being lost (Figure 4F). However, its loss did not exhibit a clear pattern, which may largely result from stochastic events during chasing. Other outcomes were rare (Figure S4A-B). It was found that intermediate drive conversion efficiency and embryo cut rare produced the most successful results (Figure 4A, C), with the lowest chasing possibility and the shortest time to population elimination. This is likely because a medium-efficiency homing suppression drive would guarantee that it would not eliminate the TARE drive, while still itself having to the efficiency needed to achieve rapid population suppression.

**Figure 4.**
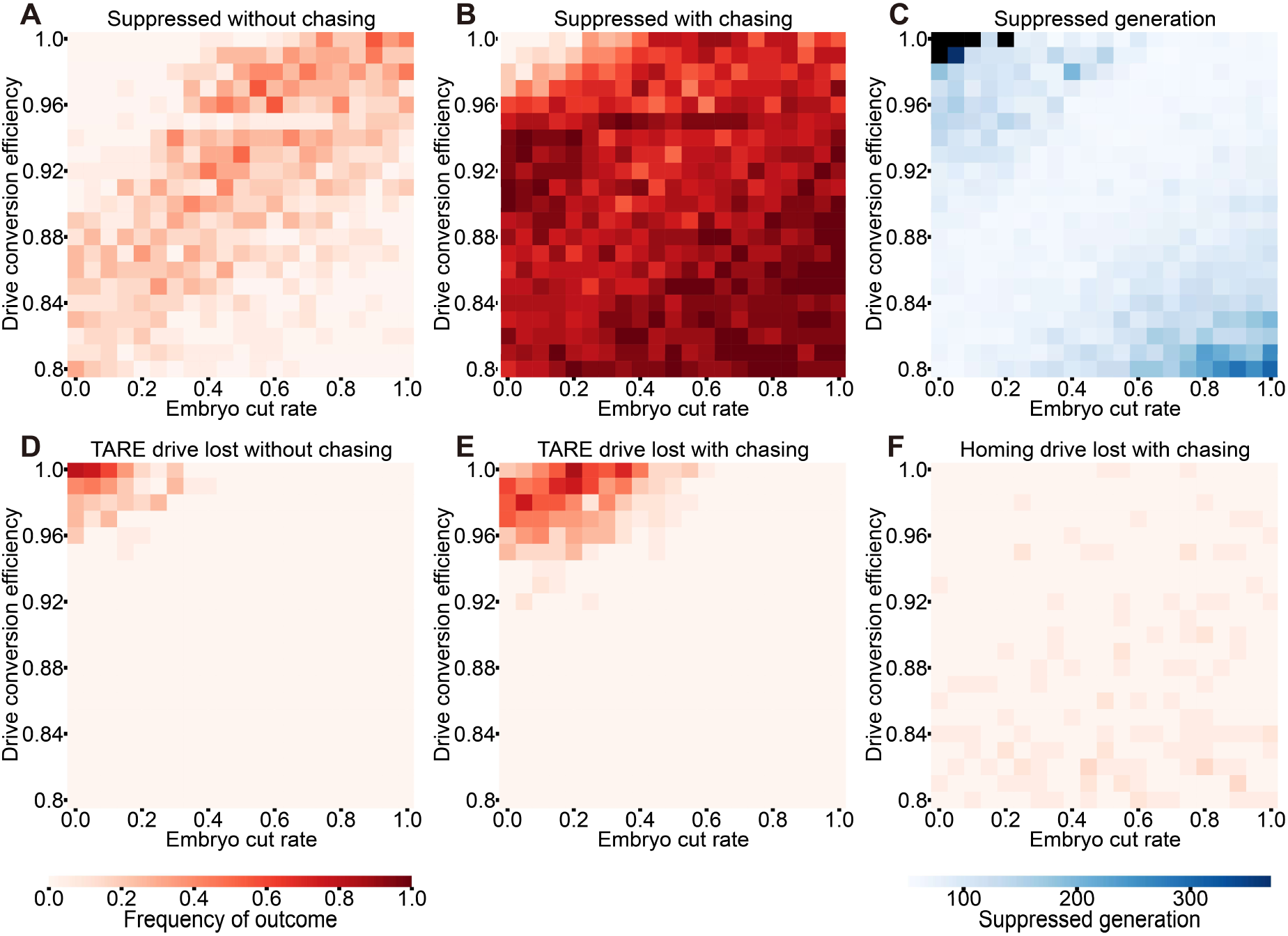
Outcomes with varying homing suppression drive germline efficiency and embryo cut rate. TARE drive homozygotes with default parameters and later homing suppression drive heterozygotes with varying germline efficiency and embryo cut rate were released from left edge to meet in the middle of a one-dimensional arena. The simulation was allowed to proceed for 1,000 generations. Outcomes with population elimination (**A)** without or (**B**) with chasing are shown. (**C**) The generation in which population elimination occurs is shown. Black squares represent parameter combinations in which population elimination did not occur. Other unsuccessful outcomes are shown, including (**D**) TARE drive loss without chasing, (**E**) TARE drive loss during chasing, and (**F**) homing drive loss during chasing. The number of simulations per spot is 20. See Figure S4 for additional rare outcomes.

While it is easier to make an effective TARE drive than a homing suppression drive, TARE drive parameters may still be suboptimal, such as small to moderate fitness costs (Champer et al., 2020b; Metzloff et al., 2022). Homing suppression drives are also known to have high fitness costs in female heterozygotes in many cases (Hammond et al., 2017; Metzloff et al., 2022; Yang et al., 2022). Varying these parameters, we found that for the TARE drive component, success is only possible when the fitness value reaches 0.94 or higher (Figure 5A-C). The mechanism of TARE drive relies on indirectly increasing its own frequency by eliminating wild-type alleles, unlike homing drives which can replicate and directly increase their frequency. Hence, the TARE drive is more sensitive to fitness costs for its general propagation ability and wave speed (Pan and Champer, 2023). As the fitness cost increased, the introduction threshold for the TARE drive also rose, preventing it from advancing in the presence of the homing drive.

**Figure 5.**
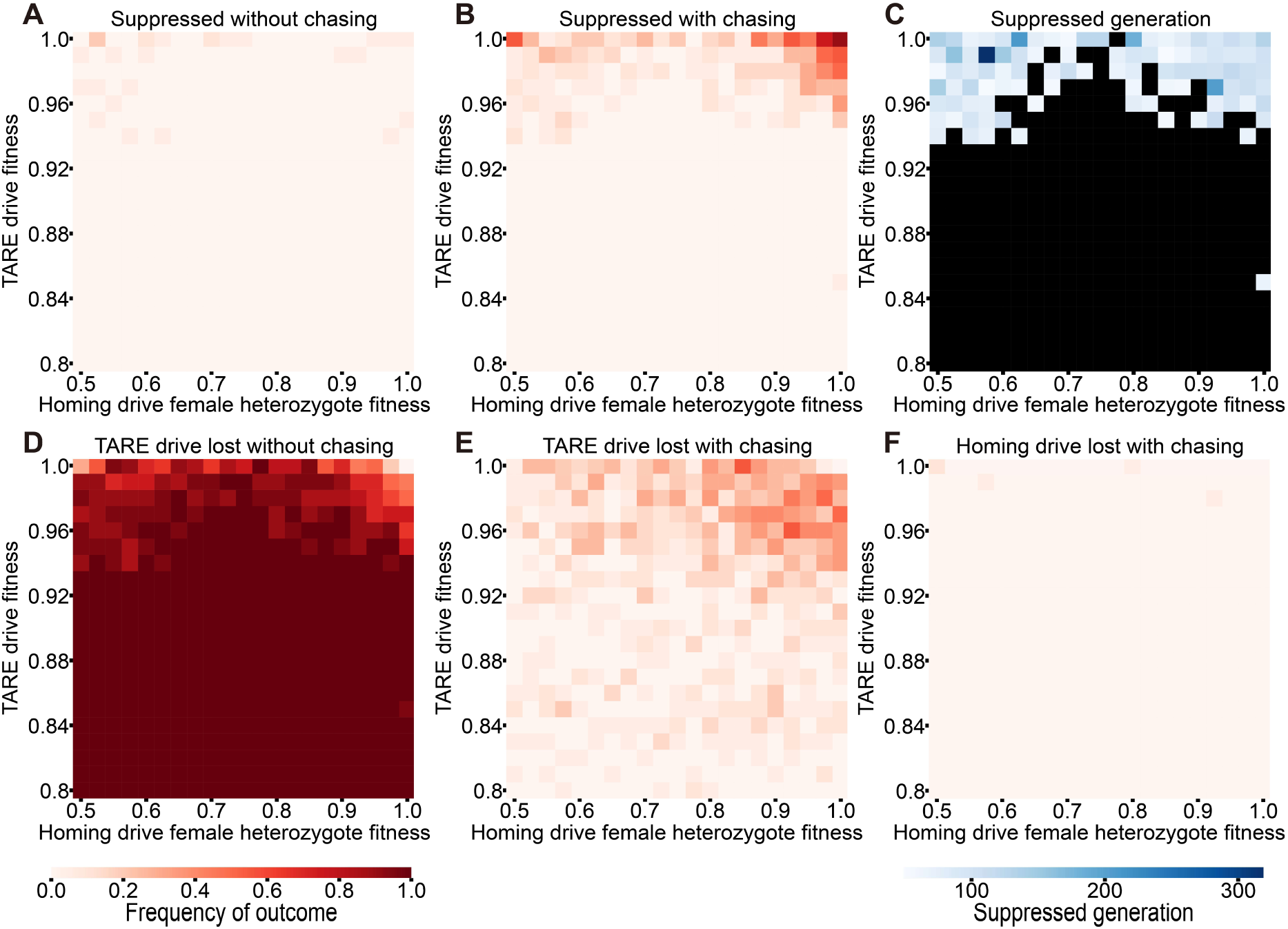
Outcomes with varying drive fitness. With default drive performance parameters and ecological parameters, TARE drive homozygotes with varying fitness and homing suppression drive heterozygotes with varying female heterozygote fitness were released from left edge with a certain time interval to meet in the middle of a one-dimensional arena. The simulation was allowed to proceed for 1,000 generations. Outcomes with population elimination (**A)** without or (**B**) with chasing are shown. (**C**) The generation in which population elimination occurs is shown. Black squares represent parameter combinations in which population elimination did not occur. Other unsuccessful outcomes are shown, including (**D**) TARE drive loss without chasing, (**E**) TARE drive loss during chasing, and (**F**) homing drive loss during chasing. The number of simulations per spot is 20. See Figure S4 for additional rare outcomes.

The homing drive component, despite having a higher tolerance for fitness costs, exhibited different tendencies across various fitness parameter ranges. Surprisingly, both higher and lower homing drive fitness levels were conducive to the success of the tethered drive system (Figure 5A-C). Those in the middle range, however, were less successful. Initially, for a TARE drive with specific fitness, when the homing drive’s fitness was low, its propagation speed was slow, allowing the TARE drive to spread more easily, which may give the whole system a better chance to succeed. As the fitness of the homing drive increased, it had a larger chance to remove the TARE drive during chasing or as stochastic loss without chasing (Figure 5D-F, S3C-D). When homing drive fitness was high, it was more stable during chasing, which also could be shorter, increasing the chance that suppression could occur.

We next assessed the effect of varying ecological parameters. Dispersal distance was varied from 0.02 to 0.08, and low-density growth rate was varied from 1.5 to 10. Successful population elimination usually occurred when the dispersal distance was high (Figure 6A-C). When dispersal distance was lower than 0.03, success was rare, perhaps because drive wave width was lower, making it easier for the suppression drive to eliminate the TARE drive (Figure 6D-E). TARE drive loss also happened when the low-density growth rate was extremely low. In this situation, once the suppression drive has reduced the population, the population’s intrinsic robustness diminishes, making it challenging to rapidly replenish population numbers, which leads to stochastic effects such as allele loss. A standard homing suppression drive could achieve a more effective suppression outcome in the situation while still suffering from higher rates of drive loss (Champer et al., 2021b). However, within the framework of the tethered homing drive for lower low-density growth rate, the TARE drive becomes more susceptible to loss earlier, leading to reduced system success. Based on the successful suppression rate and the time duration needed for suppression (Figure 6C), higher dispersal distance and intermediate low-density growth rate might be most suitable to support an effective tethered drive. Homing drive loss and low equilibrium also happen at a relatively low frequency based on rare stochastic events (Figure 6F-G, S3E), inherent in the spatial model.

**Figure 6.**
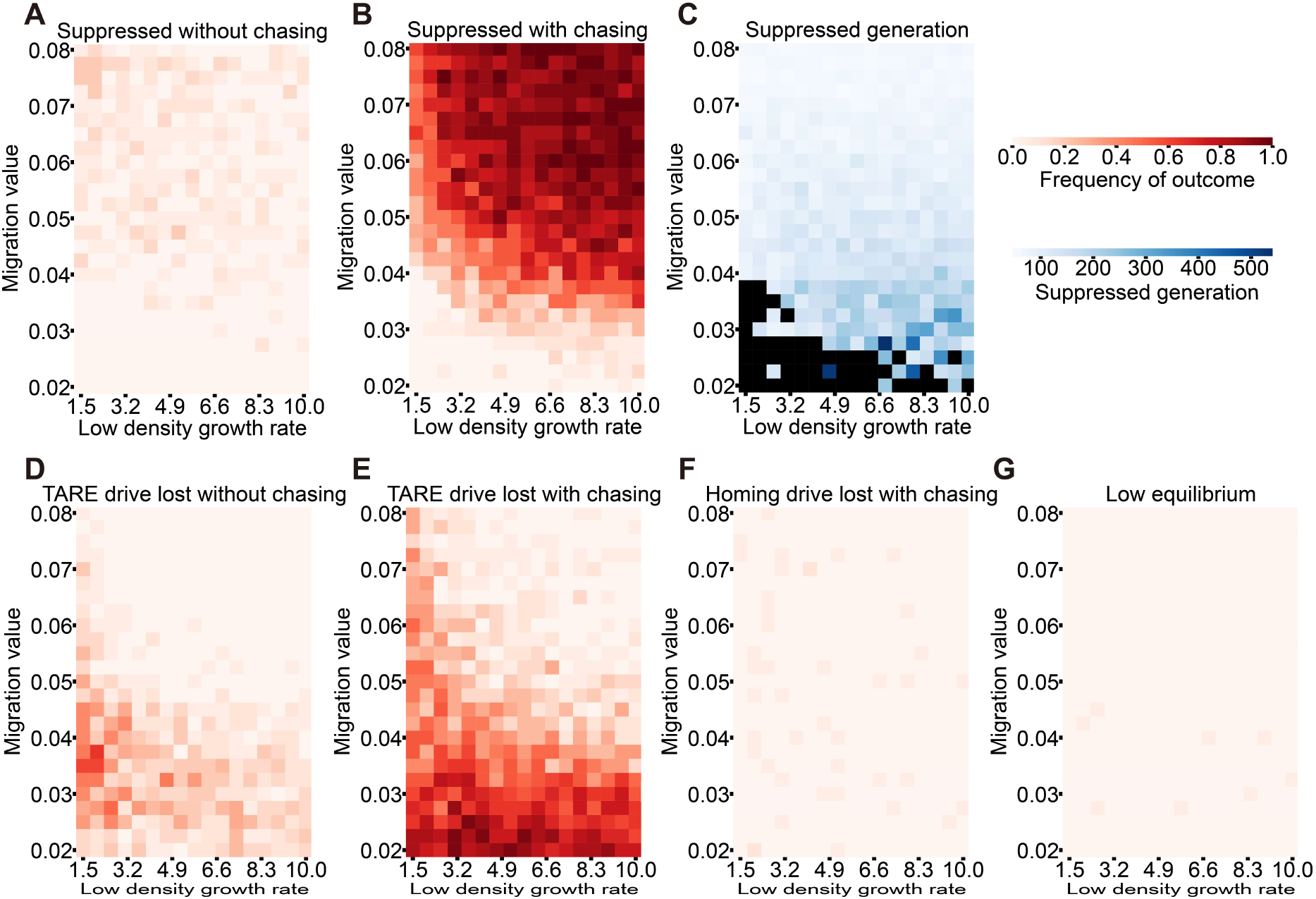
Outcomes with varying ecological parameters. TARE drive homozygotes with default parameters and homing suppression drive heterozygotes with drive conversion of 0.95 and embryo cut rate of 0.15 were released from left edge with a certain time interval to meet in the middle of one-dimensional arena and were tracked for 1,000 generations. Successful population elimination (**A**) without and (**B**) with chasing are shown. (**C**) The number of generations to suppression is shown. Black squares represent parameter combinations in which population elimination did not occur, except for the square with a dispersal distance of 0.025 and a low-density growth rate of 9.5, which occurred once after 875 generations. Other unsuccessful outcomes are shown, including (**D**) TARE drive loss without chasing, (**E**) TARE drive loss during chasing, (**F**) homing drive loss during chasing, and (**G**) low equilibrium outcome. The number of simulations per spot is 20. See Figure S4 for additional rare outcomes.

### Tethered drive outcomes in two-dimensional space

With a better understanding of wave dynamics and possible outcomes, we now extend our model from one-dimensional to two-dimensional arenas. Unlike nearly 100% success in panmictic model without default parameters, the 2D spatial models had more failure (Figure 7A). When we observe the differences between the 1D and 2D models, we find that many results of successful suppression with chasing in the 1D model become “low equilibrium” results in the 2D model, which represents a long-term, low-level coexistence of TARE drives and homing suppression drives. This indicates that chasing still occurs in two-dimensional space, but because there are more directions available for escape, wild-type individuals can manage to break free more easily from large masses of drive individuals, which then get reduced to low levels. This collapses the TARE drive wave, even though it avoids elimination. However, two dimensions still maintain the overall trend of one-dimensional models. In some parameter ranges, success it still possible, although with a reduced range. With default parameters, a moderate embryo cut rate in the homing drive can achieve better results, even though success is still not the most common outcome (Figure 7B-C).

**Figure 7.**
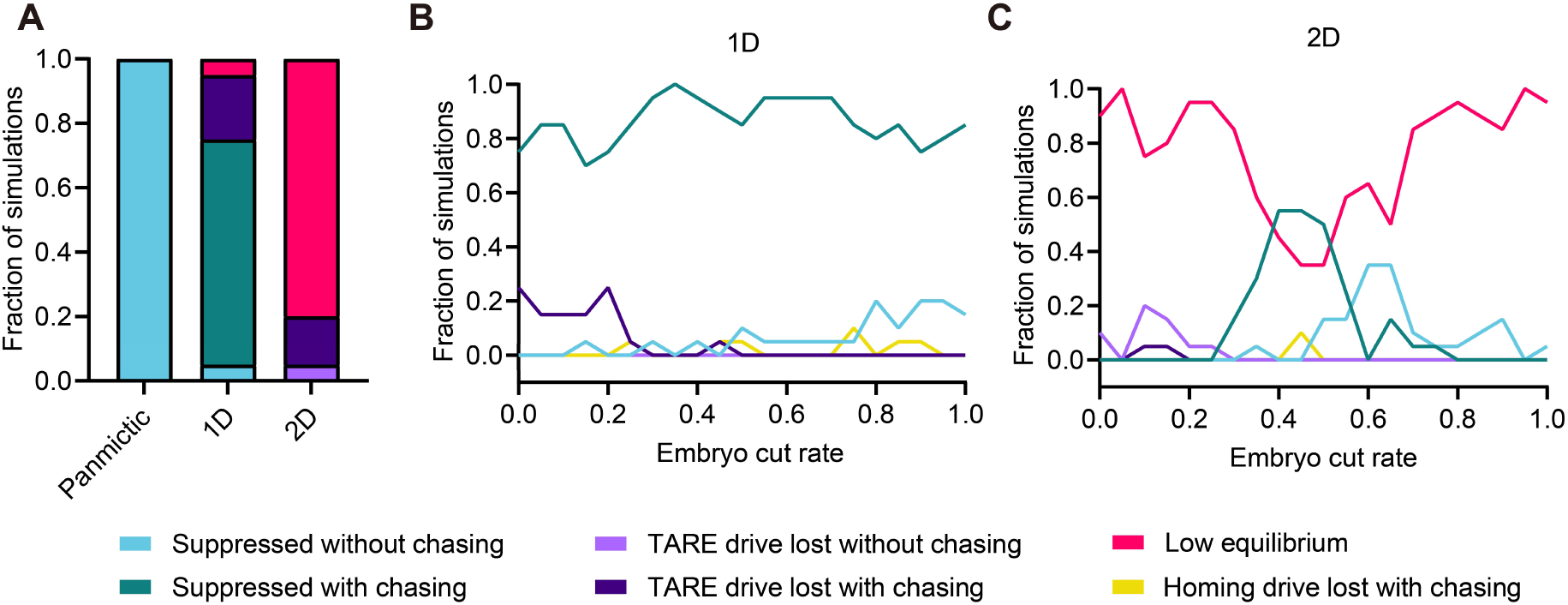
Comparison of outcomes between panmictic, 1D and 2D model with fixed parameters. (**A**) With dispersal distance of 0.05 and low-density growth rate of 5, TARE drive homozygotes with default parameters and homing suppression drive heterozygotes with germline efficiency of 0.95 and embryo cut rate of 0.15 were released and were tracked for 1,000 generations in three models, panmictic, 1D and 2D. (**B**) With varying homing drive embryo cut rate, the proportion of different simulation outcomes in 1D is shown. (**C**) The proportion of different simulation outcomes in 2D is shown. For panmictic models, the outcomes under this parameter setting are all suppressed without chasing.

Comprehensively varying homing drive conversion efficiency and embryo cut rate, the results rarely showed suppression without chasing (Figure 8A). Outcomes were primarily divided into three groups. When homing drive performance was intermediate, a thin parameter range shows mostly successful outcomes after a period of chasing (Figure 8B-C). When homing drive performance was nearly optimal, the TARE drive was quickly lost before chasing developed (Figure 8D). Most of the remaining areas are characterized by low equilibrium, indicating that in two-dimensional space, the influence of chasing and wave interference significantly increases the probability of drive failure (Figure 8E-I).

**Figure 8.**
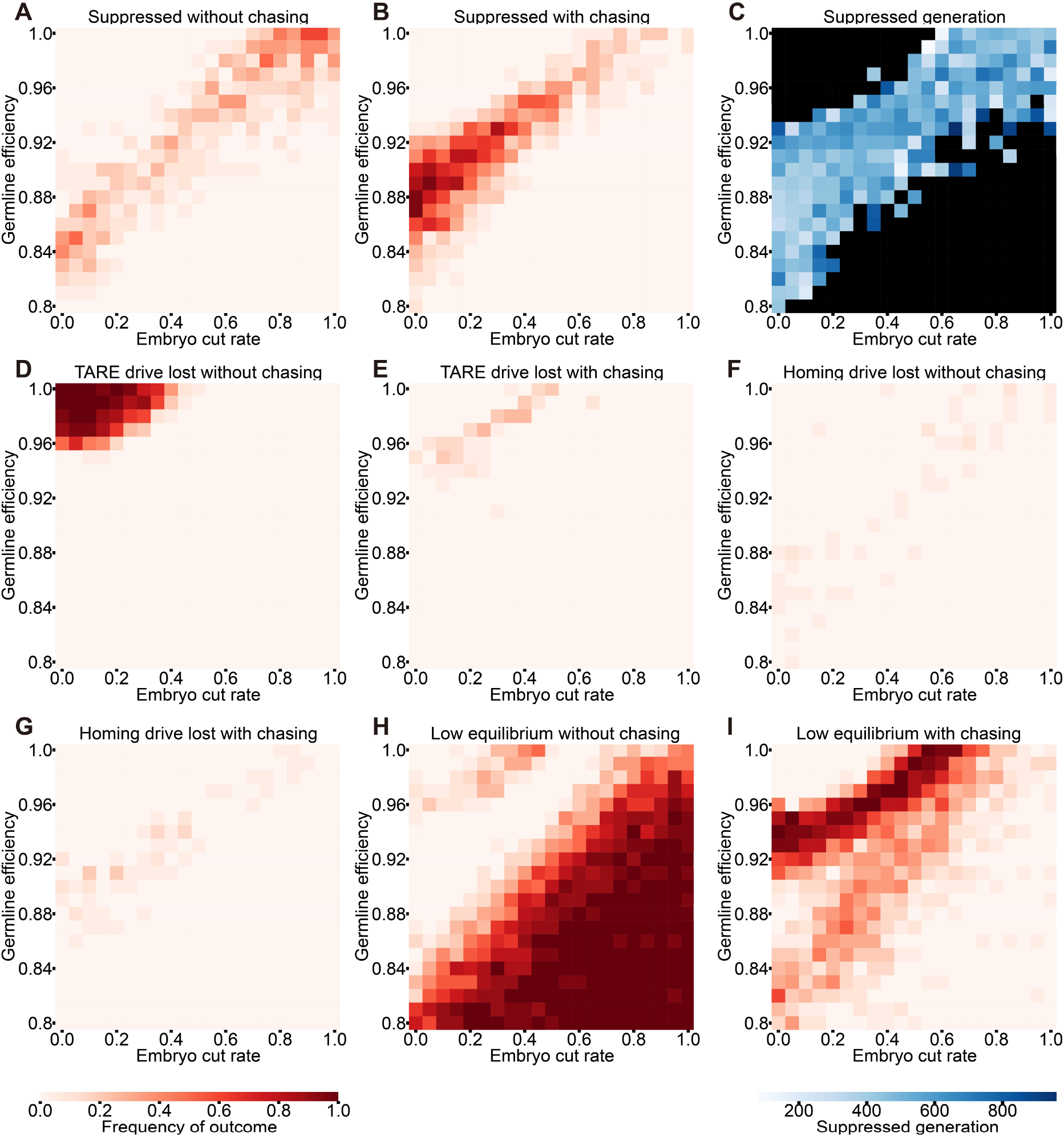
Outcomes with varying drive performance parameters in two-dimensional model. TARE drive homozygotes with default parameters and homing suppression drive heterozygotes with varying germline efficiency and embryo cut rate were released from center with a certain time interval to meet at the center of a circle, which is located in the middle of the 2D space, with a radius equal to 0.25 times the length of the space’s side and were tracked for 1,000 generations. Successful population elimination (**A**) without and (**B**) with chasing are shown. (**C**) The number of generations to suppression is shown. Black squares represent parameter combinations in which population elimination did not occur. Other unsuccessful outcomes are shown, including (**D**) TARE drive loss without chasing, (**E**) TARE drive loss during chasing, (**F**) homing drive loss without chasing, (**G**) homing drive loss during chasing, and (**H**) low equilibrium outcome without chasing, and (**I**) low equilibrium outcome after chasing. The number of simulations per spot is 20.

## Discussion

Tethered homing drives hold great promise for the genetic control of populations by combining the suppressive power of homing suppression drive and spatial restriction of confined drives like TARE drive. Yet, the greater power and speed of the suppression component can complicate the deployment of these systems. In panmictic models, depending on whether the suppression component can overcome the confined component before it reaches all individuals, tethered homing drives usually result in either successful population elimination or confined drive loss. Through the one-dimensional model, interesting interactions in tethered homing drives were shown, such as the conflict between the two components of drive. Specifically, TARE drive can reduce the relative number of wild-type, homing suppression drive can remove TARE drive and create new empty space, and wild-type can reinvade the empty space. Together, these constitute the dynamics of tethered drives in spatial models, similar to complete suppression drives in continuous space, but with more complex outcomes (Champer et al., 2021b). On one hand, considering the TARE drive’s support for the homing drive, the latter can only propagate within the areas reached by the TARE drive. On the other hand, the homing drive also becomes a brake to the TARE drive, affecting its activity. With such a constraining relationship, it was shown that the best results could obtained with a perfect tethered drive and imperfect homing drive, which might also be closer to the reality of current technology.

The tolerance for the drive performance parameters of the TARE drive’s fitness and the exact range of homing drive imperfections is more limited, with only a relatively narrow parameter range facilitating favorable drive outcomes. This serves as a caution for the future design, emphasizing the need to pay close attention to fitness costs and embryo resistance levels. Similarly, ecological parameters significantly influence drive outcomes, suggesting that when we apply the tethered homing drive to different populations with different ecological characteristics, we should tailor our strategy. However, actual dispersal could be quite a bit higher than we modeled, expanding the possible success range when varying other parameters. To avoid failure from simultaneous or closely timed releases, we can also simply adopt a prudent two-step release strategy in most real-world scenarios: first, wait for the TARE drive to nearly complete its spread, and then release the homing suppression drive far enough from the TARE drive boundary that it has no chance to catch up before the TARE drive reaches the whole population. Though this may involve some delays and need for coordination, it would also remove complexity of drive interaction from the scenario.

So far, modification drive designs have been relatively more diverse and flexible than suppression drives, particularly for confinement. As a flexible and controllable two-step method, tethered drive passes on advantages of confined modification drives to a suppression drive. As one of the major categories of modification drive, the CRISPR toxin-antidote drive family has large numbers of variants, with different introduction thresholds and corresponding confinement levels (Champer et al., 2021a; Champer et al., 2020a). Some toxin-antidote variants even have a drive-performance dependent threshold. This confinement allows use of strong homing suppression drives in a greater variety of scenarios. However, we must not overlook the fact that when we create these linked systems with two parts, they may exhibit different properties. In a single-construct confined suppression drive system such as TADE suppression, although its spread is reduced by the suppression component, drive loss is usually avoided because the system is integrated. In contrast, the suppression part of a tethered drive can be much faster than the confined part, so it can catch up and cause interference. Nonetheless, such a system can overall still be faster than a single-construct system, depending on the exact parameters for both the drive and ecology of the system.

For potential practical application, we also proposed general release plans for either panmictic populations or a widespread release. If this simultaneous release plan is adopted, we should make sure that the release ratio of TARE drive to homing suppression drive is sufficiently high. Usually, a small homing drive release sufficient to avoid stochastic loss coupled with a TARE drive release somewhat above its normal threshold would be sufficient for success. Additionally, we also need to carefully consider how to mitigate unforeseen failures should they occur. If the initial release design can meet appropriate parameter ranges to ensure efficacy, it is naturally most robust to unexpected events. However, a worst-case scenario of both drives persisting at low levels occurs, a post hoc release is necessary to break the equilibrium, though this was successful in less than 40% of cases when it took place in only a fraction of the total area. If a wider release in this case could be approximated by a panmictic model, then there should be a sufficiently high level that could ensure success, though this may not be economically feasible in some cases. In most cases, the supplemental release will not lead to suppression, but will at least eliminate TARE drive (Figure S5), which would eventually be followed by the homing drive, resetting the population. This is similar to a “brake-driven gene drive reversal” strategy, aiming to stop the spatial spread of a drive and restore the population (Girardin et al., 2019; Vella et al., 2017; Wu et al., 2016). From this perspective, the dynamic process between the homing suppression drive and the TARE drive is also a “catch me if you can” scenario, in which the goal is to use a second drive to eliminate the first drive. In this case, “success” involves TARE drive loss. Because elimination of the TARE drive by the homing drive is possible over a much wide range of parameter than elimination of another homing drive, it could be considered as a promising backup intervention for a modification strategy, aligning with the concept of “socially responsible drives” (Tanaka et al., 2017).

To gain greater understanding of tethered drives, models were extended from panmictic to one-dimensional to two-dimensional space. We can clearly see the significant differences in results between each of these. The panmictic model typically produces a binary outcome; under fixed parameters, it either succeeds or fails, with very little uncertainty in the parameter space. In contrast, the spatial model exhibits a complex distribution of results. In spatial environments, there are often greater uncertainties caused by founder events and subsequently greater genetic drift (Slatkin and Excoffier, 2012), accounting for some of the variation we see.

Both one-dimensional and two-dimensional spaces have their distinct and different advantages. In one-dimensional simulations, we can more intuitively observe the chasing process between the homing suppression drive, TARE drive, and wild-type, as well as the diverse outcomes produced when waves formed by individuals come into contact. Additionally, there is a clearer definition of penetration and escape. Such models are also more similar to certain corridor and hub areas in real populations. However, expanding from one dimension to two can better simulate more real-world scenarios. In two-dimensional space, individual movements are more varied. Thus, our two-dimensional simulation results may have similar types of outcomes as one-dimensional simulations, but also further restrict the parameters range for success.

Tethered drive systems offer promising avenues for genetic control of populations by combining the suppressive power of homing suppression drives with the potential different levels of spatial restriction offered by confined drives such as toxin-antidote drives. This system demonstrates complex interactions between its confined and suppression parts, highlighting the interplay between drive and wild-type, as well as the dynamics across different genotypes. Expanding from panmictic models to spatial models reveals greater complexity in outcomes, underscoring the need for precise parameter selection or careful release planning to ensure successful application.

## Acknowledgements

The cluster-based data collection was assisted by High-Performance Computing Platform of the Center for Life Science at Peking University. This study was supported by the Center for Life Sciences and the National Science Foundation of China (grant 32270672).

## Supplemental Information

**Figure S1.**
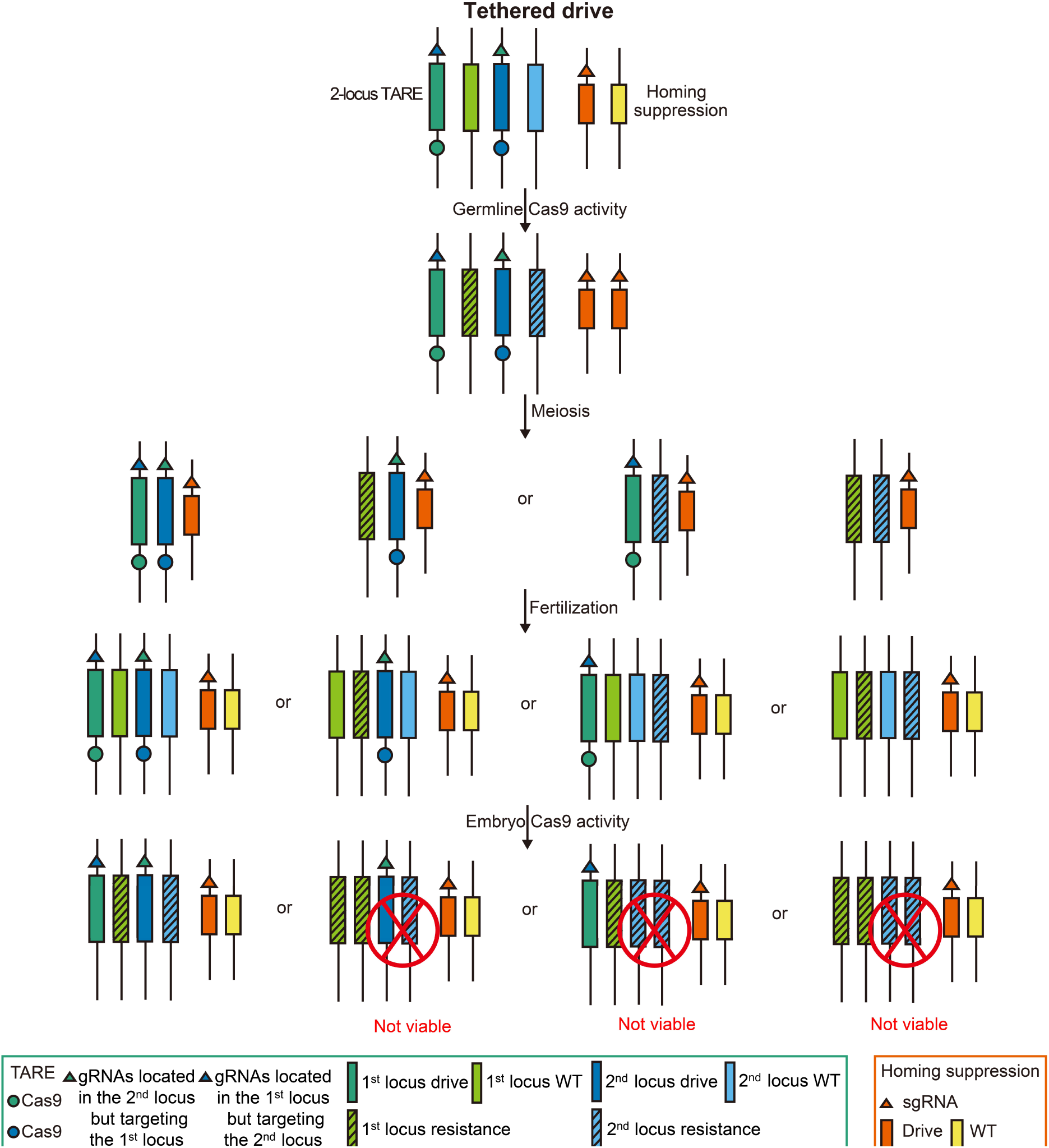
2-locus TARE-modified tethered homing drive mechanism. The mechanism of drive activity is shown in an example of an individual that is homozygotes for 2-locus TARE drive and heterozygotes for homing suppression drive mating with a wild-type individual.

**Figure S2.**
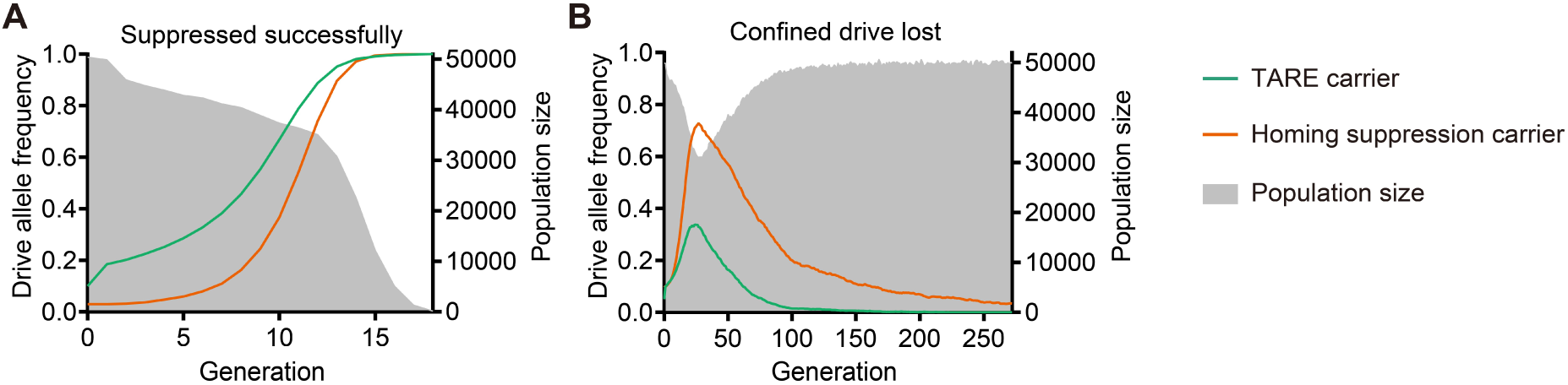
Primary outcomes in tethered homing drives in panmictic model. Time-series of different possible outcomes is shown. Green and orange solid lines represents the drive allele frequency of TARE drive and homing suppression drive, respectively. The grey shadow represents the population size. (**A**) An example with successful population elimination is shown. (**B**) An scenario with confined drive loss is shown.

**Figure S3.**
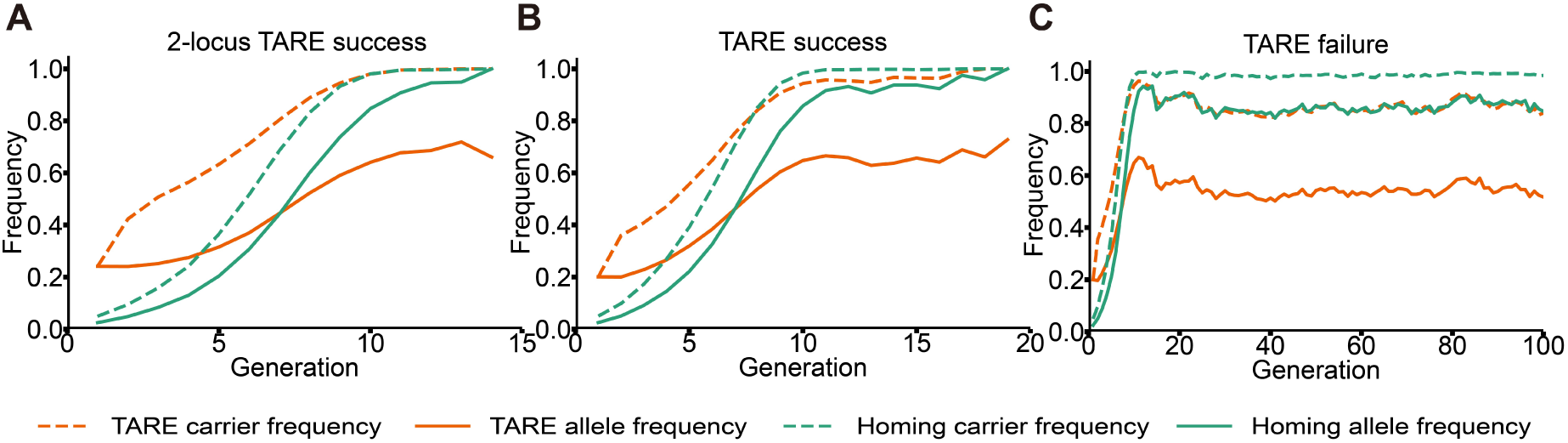
Rare high equilibrium outcome in the panmictic model. Time-series of different possible outcomes is shown. (**A**) The typical frequency trajectory of the 2-locus TARE drive in a tethered drive suppression scenario is shown. (**B-C**) Two different frequency trajectories for the TARE drive scenario. Note that that homing suppression drive carrier frequency increases above the TARE drive in both these cases, but due to stochastic events, the TARE drive carrier frequency is not able to increasing to 100% in the failure scenario, dropping to a lower equilibrium frequency.

**Figure S4.**
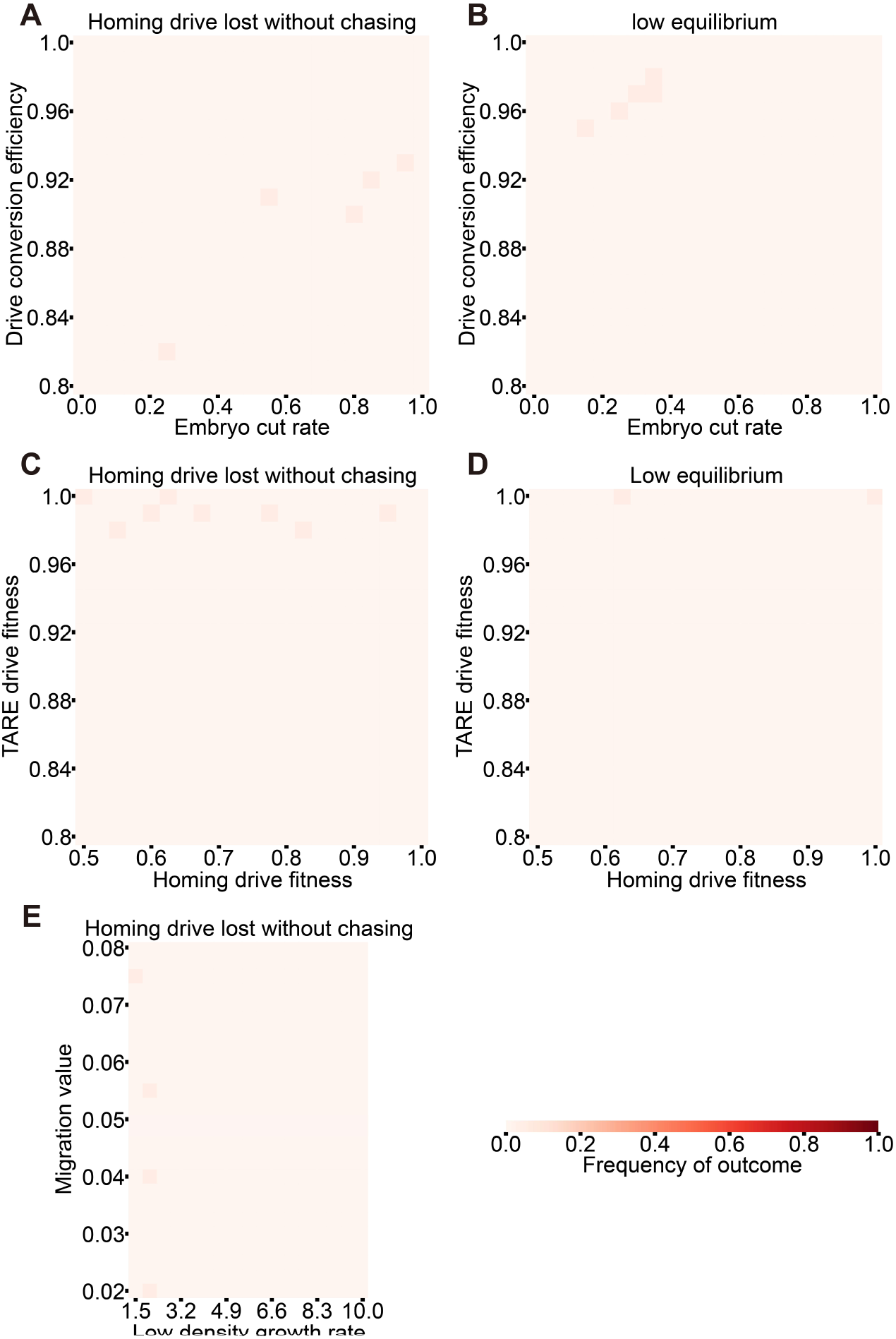
Infrequent outcomes with varying parameters in one-dimensional models. (**A-B**) Additional outcomes in systems with varying homing suppression drive embryo cut rate and drive conversion rate are shown. (**C-D**) Additional outcomes in systems with varying TARE fitness and homing suppression drive somatic fitness are shown. (**E**) An additional outcome in drives with varying migration rate and low-density growth rate is shown.

**Figure S5.**
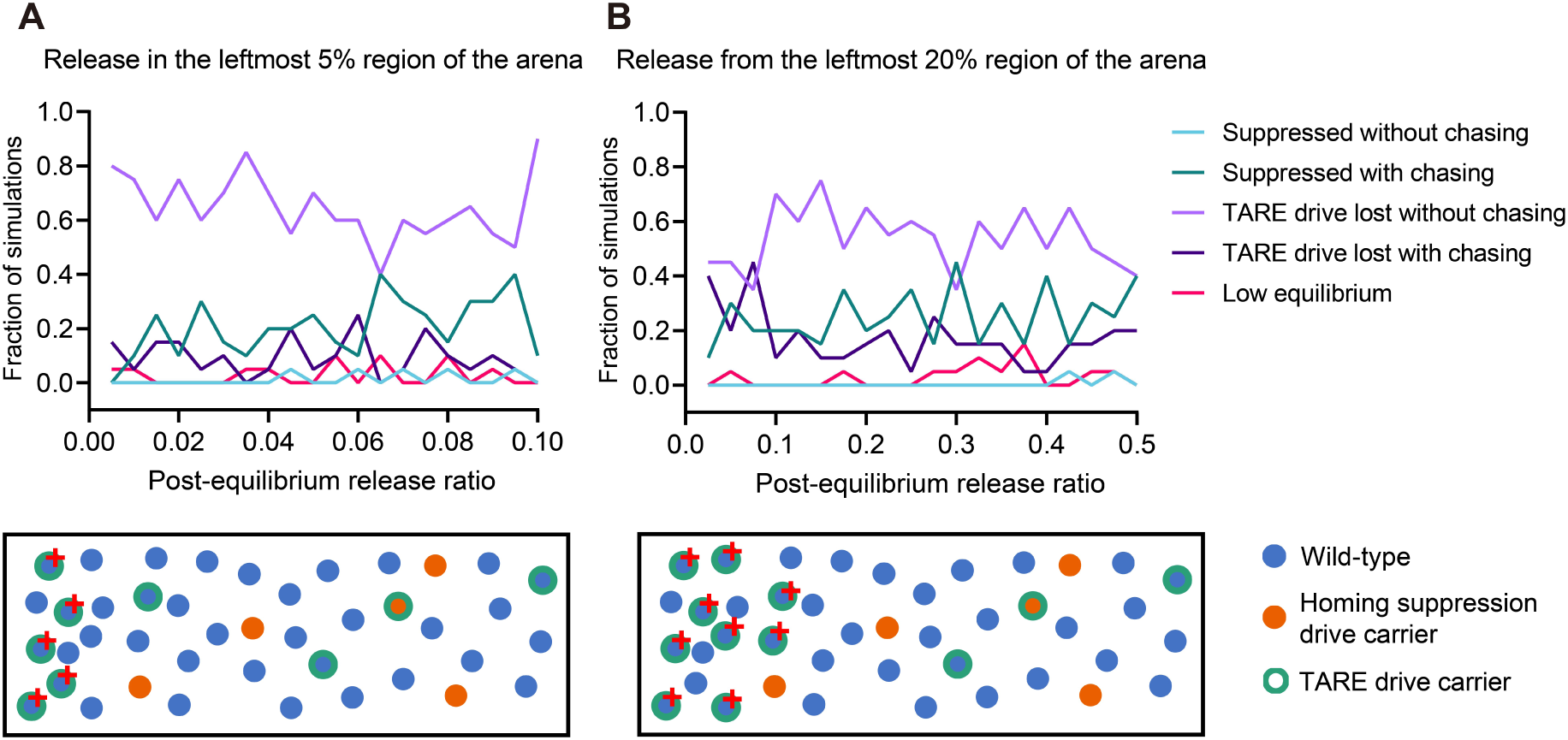
A remedial release after low equilibrium outcome. After a low equilibrium outcome in the 1D scenario, new TARE drive individuals are released from the left edge at the post-equilibrium release ratio (the relative level compared to the whole population). Major outcomes are shown in different colors in the upper area. The corresponding release schematics are shown in the lower area, and the population dynamics were tracked for 1,000 generations. In the release schematic, individuals marked with a red cross symbolize newly released individuals intended for remediation. (**A**) New individuals are released from the leftmost 5% region of the arena with a proportion ranging from 0.5% to 10% of the whole population. (**B**) New individuals are released from the leftmost 20% region of the arena with a proportion ranging from 0.25% to 50% of the whole population.

